# The Genetic Basis of Differential Autodiploidization in Evolving Yeast Populations

**DOI:** 10.1101/2021.03.10.434832

**Authors:** Sudipta Tung, Christopher W. Bakerlee, Angela M. Phillips, Alex N. Nguyen Ba, Michael M. Desai

**Author notes:** These authors contributed equally to this work and are joint first authors (Sudipta Tung, Christopher W. Bakerlee). Department of Biology, Ashoka University, Sonipat, Haryana 131029, India (Sudipta Tung). Department of Cell and Systems Biology, University of Toronto, Toronto, Canada (Alex N. Nguyen Ba).

## Abstract

Spontaneous whole-genome duplication, or autodiploidization, is a common route to adaptation in experimental evolution of haploid budding yeast populations. The rate at which autodiploids fix in these populations appears to vary across strain backgrounds, but the genetic basis of these differences remains poorly characterized. Here we show that the frequency of autodiploidization differs dramatically between two closely related laboratory strains of *Saccharomyces cerevisiae*, BY4741 and W303. To investigate the genetic basis of this difference, we crossed these strains to generate hundreds of unique F1 segregants and tested the tendency of each segregant to autodiplodize across hundreds of generations of laboratory evolution. We find that variants in the *SSD1* gene are the primary genetic determinant of differences in autodiploidization. We then used multiple laboratory and wild strains of *S. cerevisiae* to show that clonal populations of strains with a functional copy of *SSD1* autodiploidize more frequently in evolution experiments, while knocking out this gene or replacing it with the W303 allele reduces autodiploidization propensity across all genetic backgrounds tested. These results suggest a potential strategy for modifying rates of spontaneous whole-genome duplications in laboratory evolution experiments in haploid budding yeast. They may also have relevance to other settings in which eukaryotic genome stability plays an important role, such as biomanufacturing and the treatment of pathogenic fungal diseases and cancers.

## INTRODUCTION

As populations evolve, they occasionally undergo changes in ploidy. These changes have led to extensive ploidy variation across the tree of life, including notable differences among fungi (Albertin and Marullo 2012), plants, animals, and other eukaryotes (reviewed in Otto 2007; Sémon and Wolfe 2007). Ploidy changes and broader genome instability have also been observed in clinically relevant contexts, where they appear to contribute to fungal pathogenesis (Gerstein et al. 2017) and tumorigenesis (Fujiwara et al. 2005; Storchova and Kuffer 2008).

In several recent laboratory evolution experiments with *Saccharomyces cerevisiae*, populations have been found to spontaneously duplicate their whole genomes, or autodiploidize, with high frequency in the early stages of adaptation (Fisher et al. 2018; Gerstein et al. 2006; Gorter et al. 2017; Kosheleva and Desai 2018; Levy et al. 2015; Nguyen Ba et al. 2019; Voordeckers et al. 2015; Hong and Gresham 2014; Oud et al. 2013). In one such experiment, autodiploidization events were found to have a substantial fitness benefit, and make up the vast majority of initial beneficial mutations (Venkataram et al. 2016). Autodiploidization occurs in these strains despite mutations at the homothallic switching endonuclease (*HO*) locus that sharply reduce the frequency of mating-type switching (Haber et al. 1980).

While some work has been done to illuminate how different environmental conditions affect the propensity for autodiploids to arise and increase to appreciable frequency (Harari et al. 2018), the genetic basis of this trait remains uncharacterized. This leaves a significant gap in our understanding of perhaps the most commonly observed mutation in yeast laboratory evolution experiments. This gap also presents a practical challenge for researchers conducting yeast evolution experiments, where autodiploidization frustrates efforts to study the evolutionary consequences of ploidy-dependent population genetic parameters, including mutation rates, recombination, and the distribution of fitness effects. In addition, autodiploidization can complicate efforts to genetically manipulate budding yeast, such as by adding DNA barcodes (Levy et al. 2015) or activating more complex genetic circuitry (e.g., Cre-Lox recombination machinery (Nguyen Ba et al. 2019)), especially in the context of long-term culture. Thus, a better understanding of the genetic basis of this trait may benefit both researchers in experimental evolution and those who use or study yeast in industry, medicine, and molecular biology.

Previous evolution experiments founded with haploid clones derived from budding yeast strains BY4741 and W303 have suggested that BY-derived populations fix autodiploids more frequently than W303-derived populations (e.g., (Levy et al. 2015; Gorter et al. 2017; Voordeckers et al. 2015; Hong and Gresham 2014 for BY; Johnson et al. 2021; Jerison et al. 2020 for W303, but see Fisher et al. 2018)). Here, we combine experimental evolution with a QTL mapping approach to identify the genetic basis for this difference in propensity to autodiploidize. Consistent with recent work describing the genetic basis for aneuploidy tolerance in wild yeast (Hose et al. 2020, although see Scopel et al. 2020) we identified alleles of the *SSD1* gene as the primary genetic determinant of this difference. Below, we describe the experiments that led to this finding and its confirmation, and we speculate briefly about the underlying biological mechanism.

## MATERIALS AND METHODS

### Yeast strains and F1 segregants for QTL mapping

To generate F1 segregants for QTL mapping, we used BY-derived YAN463 (*MAT**a**, his3Δ1, ura3Δ0, leu2Δ0, lys2Δ0, RME1pr::ins-308A,ycr043cΔ0::NatMX,ybr209w::CORE-UK, can1::STE2pr_SpHIS5_STE3pr_LEU2*) as the parent that frequently autodiploidized, while W303-derived yGIL646 (*MAT**α**, ade2-1, CAN1, his3-11, leu2-3,112, trp1-1, bar1Δ::ADE2, hmlαΔ::LEU2, GPA1::NatMX, ura3Δ::PFUS1-yEVenus*), described elsewhere (Fisher et al. 2018), served as the parent that rarely autodiploidized (Figure S1). Note that we included the *RME1pr::ins-308A* mutation in our BY strain to increase its sporulation efficiency. The *CORE-UK* cassette was originally included to facilitate knocking new genetic material into the *YBR209W* locus via the *delitto perfetto* method (Storici and Resnick 2006), but it was incidental to this study. After mating and sporulation, we isolated a total of 627 haploid F1 offspring (segregants), in three separate sets. The first set of segregants was constructed by dissecting 65 tetrads, yielding 260 “tetrad spores.” The second and third sets consisted of 184 and 183 *MAT***a** F1 segregants respectively, each set with common auxotrophies, which we isolated by germinating spores on synthetic defined (SD) growth medium with canavanine but without adenine, histidine (SD –Ade –His +Can), and without adenine, histidine, uracil, tryptophan (SD –Ade –His –Ura –Trp +Can), respectively. Note that since the W303 strain was auxotrophic for histidine and BY’s *Schizosaccharomyces pombe*-derived *HIS5* (orthologous to *S. cerevisiae’s HIS3*) was under control of the *MAT***a**-specific *STE2* promoter, we were able to select for *MAT***a** spores by excluding histidine from the selection media. We refer to these segregant sets as “selected spores” hereafter.

### Experimental evolution

To assess autodiploidization propensity, we founded seven replicate populations from individual clones of each of the two parental genotypes, and one replicate population from each of the 627 F1 segregants. We propagated each of the resulting 641 populations for 500 generations in unshaken flat bottom polypropylene 96-well microplates using a standard batch culture protocol (with 1:2^10^ dilutions every 24 hours). All evolution was conducted at 30°C in 128μL of YPD (a rich laboratory media; 1% Bacto yeast extract (VWR #90000– 722), 2% Bacto peptone (VWR #90000–368), 2% dextrose (VWR #90000–904)) with ampicillin sodium salt (100ug/mL (VWR #97061-440)). All liquid handling was conducted using a BiomekFX robot (Beckman Coulter), as described previously (see e.g. Lang et al. 2011). To detect contamination and cross-contamination events, each 96-well plate contained a unique pattern of “blank” wells containing only media. No contamination was observed in the blank wells at any point during this experiment. At 50-generation intervals, we froze aliquots of all populations in 10% glycerol at −80°C. Prior to conducting ploidy assays and sequencing library preparation, we revived the relevant populations by thawing and inoculating 4 μL of each into 124 μL YPD at 30°C.

### Examining ploidy by nucleic acid staining

After evolving for 500 generations, we evaluated the ploidy status of each population by staining for DNA content using a procedure previously described (Jerison et al. 2020; Johnson et al. 2020), with slight modifications. Briefly, 6μL of saturated culture from each population was added to 120μL water in a fresh 96-well plate and centrifuged (2,000 rcf, 2 minutes). To fix the cells, supernatants were removed, and the pellets were resuspended by gentle pipetting in 150μL of 70% ethanol and incubated for 1h at room temperature. The samples were then centrifuged (2,000 rcf, 2 minutes), supernatants were removed, and cells were resuspended in 65μL RNAase A solution consisting of 10 mg/mL RNAase A (VWR Life Science, 9001-99-4) dissolved in 10 mM Tris-HCl, pH 8.0 and 15 mM NaCl, and incubated for ~4h at 37°C. Subsequently, 65μL of 2μM SYTOX green (Thermo Fisher Scientific, S7020) in 10 mM Tris-HCl, pH 8.0 was added to each sample, shaken briefly on a Titramax 100 plate shaker (Heidolph Instruments) for approximately 30 seconds, and incubated in the dark for at least 20 minutes at room temperature. The samples were then analyzed using a Fortessa flow cytometer (BD Biosciences). DNA content of ~ 10,000 cells of each sample was measured through a linear FITC channel and, using Flowing software version 2.5.1 (Turku Bioscience), FITC histograms (Figure S2, S3 and S7) were compared to known haploid and diploid controls to identify their ploidy.

### Genotyping with whole-genome sequencing

We genotyped all 260 F1 segregants from the tetrad spore set using whole-genome Illumina sequencing at ~5X coverage, and the parental strains YAN463 and yGIL646 at 125X and 40X coverage, respectively. To account for parental differences in auxotrophies at lysine and tryptophan, which we suspected might affect autodiploidization propensity, we grouped “selected spores” based on their lysine and tryptophan auxotrophy and ploidy status after evolution and sequenced the eight resulting pooled samples (Lys proto-/auxotrophy × Trp proto-/auxotrophy × haploid/autodiploidized).

To prepare sequencing libraries for all samples in parallel, we used a BiomekFXP liquid handling robot (Beckman Coulter) to extract total genomic DNA from ~500 μL saturated cultures of all samples, following a previously described procedure (Johnson et al. 2020). A high-throughput Bio-On-Magnetic-Beads (BOMB) protocol with paramagnetic beads and GITC lysis buffer (Oberacker et al. 2019) was used for this step, followed by DNA quantification using the AccuGreen™ High Sensitivity dsDNA Quantitation kit (Biotium, 31066) on clear flat-bottom 96-well polystyrene plates (Corning®, VWR Life Science, 25381-056). Extracted genomic DNA was then subjected to Nextera tagmentation (following Baym et al. 2015) in preparation for multiplexed Illumina sequencing. Tagmented PCR products were then purified using PCRcleanDX magnetic beads (Aline Biosciences) through a two-sided size selection procedure with 0.5/0.75X or 0.5/0.8X bead buffer ratios (Johnson et al. 2020). Quality of the multiplexed libraries was verified by estimating their fragment-size distributions using the Agilent 4200 TapeStation system and sequenced with 2×150bp paired-end chemistry on Illumina NextSeq 500 and Illumina NovaSeq platforms.

After obtaining raw sequence reads, we first trimmed them using `Trimmomatic` v0.36 (Bolger et al. 2014). We then aimed to obtain parental reference genomes and construct a list of the single nucleotide polymorphisms (SNPs) that are different between them. First, we subjected the reads for the BY-derived parent, YAN463, to a `Breseq` v0.31.0 pipeline (Deatherage and Barrick 2014) with BY4742 genome assembly reference sequence (GCA_003086655.1) in order to identify variants. Using `Breseq’s gdtools` utility program, the identified variants were applied back into the BY4742 reference genome to create an updated BY-parental genome reference. Next, the reads for the W303-derived parent, yGIL646, were parsed through `Breseq` v0.31.0 pipeline using the newly constructed updated BY-parental genome as a reference. The identified SNPs were incorporated into the updated BY-parental genome reference using `Breseq’s gdtools` utility program to construct an updated W303-parental genome reference. This ensured that the location of each SNP is identical in both parental genome references. The parental genome references were then compared to identify a list of 8505 SNPs, differing between these two genome references. Subsequently, this list of SNPs was used to identify from which parent (BY or W303) each locus was inherited in all the tetrad spores. In short, sequences for each tetrad segregant were checked for appropriate coverage and quality, the reads were aligned to BY- and W303-parental reference genome sequence separately using `bowtie2` and indexed using `samtools`. We identified the number of reads matching each parental reference at each locus using `Python` and inferred genotype at each of these loci using a hidden Markov model (HMM) algorithm. Sequences for two segregants were disregarded due to insufficient read count.

Similarly, for the eight pooled samples of the selected spores, the number of sequencing reads matching BY and W303 parental sequences at each of the 8505 loci was computed. This data was used for the enrichment analysis described below.

### QTL mapping

Our dataset consisting of genotypes (B or W, corresponding to the BY and W303 parental background respectively) at 8505 loci (columns) of 258 segregants (rows) and their ploidy phenotype after evolution (binary data, haploid = 0, diploid = 1) was used as the input for QTL analysis using R/qtl v1.46-2 software as described below (following Broman and Sen 2009). Before QTL mapping, a battery of diagnostic probes, involving a test for segregation distortion of the markers and an analysis of anomalous genotyping similarity and number of crossover events for the segregants, were checked to avoid spurious mapping (see supplementary Text S1 for details). This resulted in a clean dataset consisting of genotypes of 255 segregants at 8475 loci with their corresponding phenotypes, which then entered the following QTL mapping pipeline.

First, we computed LOD scores for all 8475 loci assuming the presence of a single QTL using standard interval mapping and the Haley–Knott regression method for a binary phenotype with LOD significance thresholds computed from 10000 permutations. Next, to find any potential interactions between multiple QTLs, we divided our data into predictor and test datasets. We chose 150 segregants arbitrarily to form a predictor dataset and subjected their genotype and phenotype data to a forward/backward stepwise search algorithm (`stepwiseqtl`) with LOD significance thresholds computed from 1000 permutations. Based on the LOD score profile of single-QTL analysis above (see Results) we restricted this search to chromosomes IV and XIV only, and the maximum number of QTLs allowed in a model was kept to 4. Subsequently, we fitted the predicted QTL model onto the remaining data consisting of 105 segregants (test dataset) using `fitqtl` followed by the `refineqtl` function.

Further, to reveal any additional low-effect QTL for the autodiploidization phenotype, we rescanned the data using single-QTL analysis methods after regressing out the QTL with highest LOD score obtained above. Effect sizes of the two alleles of the QTL with statistically significant LOD score were estimated using the `effectplot` function.

### Enrichment analysis

For each of the eight pooled samples (Lys proto-/auxotrophy × Trp proto-/auxotrophy × haploid/autodiploidized) of ‘selected spores,’ we scanned their sequencing reads at the SNP that led to statistical significance in the QTL analysis above. The proportion of those reads matching with BY version of the QTL locus was computed to find whether this statistic was different in the haploid and diploid pool.

### Experimental validation of QTL mapping result

To validate the results of our QTL mapping analysis, we cleanly knocked out the entire open reading frame (ORF) of the gene containing the statistically significant QTL, *SSD1*, using a HygMX or KanMX cassette in BY4741, W303-derived yGIL104, and RM-derived YAN516 (Table 1). Our HygMX cassette, which conferred resistance to hygromycin, was under *Ashbya gossypii TEF1* promotion and termination (Wach et al. 1994). Our KanMX cassette, which conferred resistance to G418, was under control of the *TEF1* promoter from *Kluyveromyces lactis* and under tSynth8 termination (Curran et al. 2015). The KanMX-constructed strains were used in the subsequent lab evolution experiment.

**Table 1:**
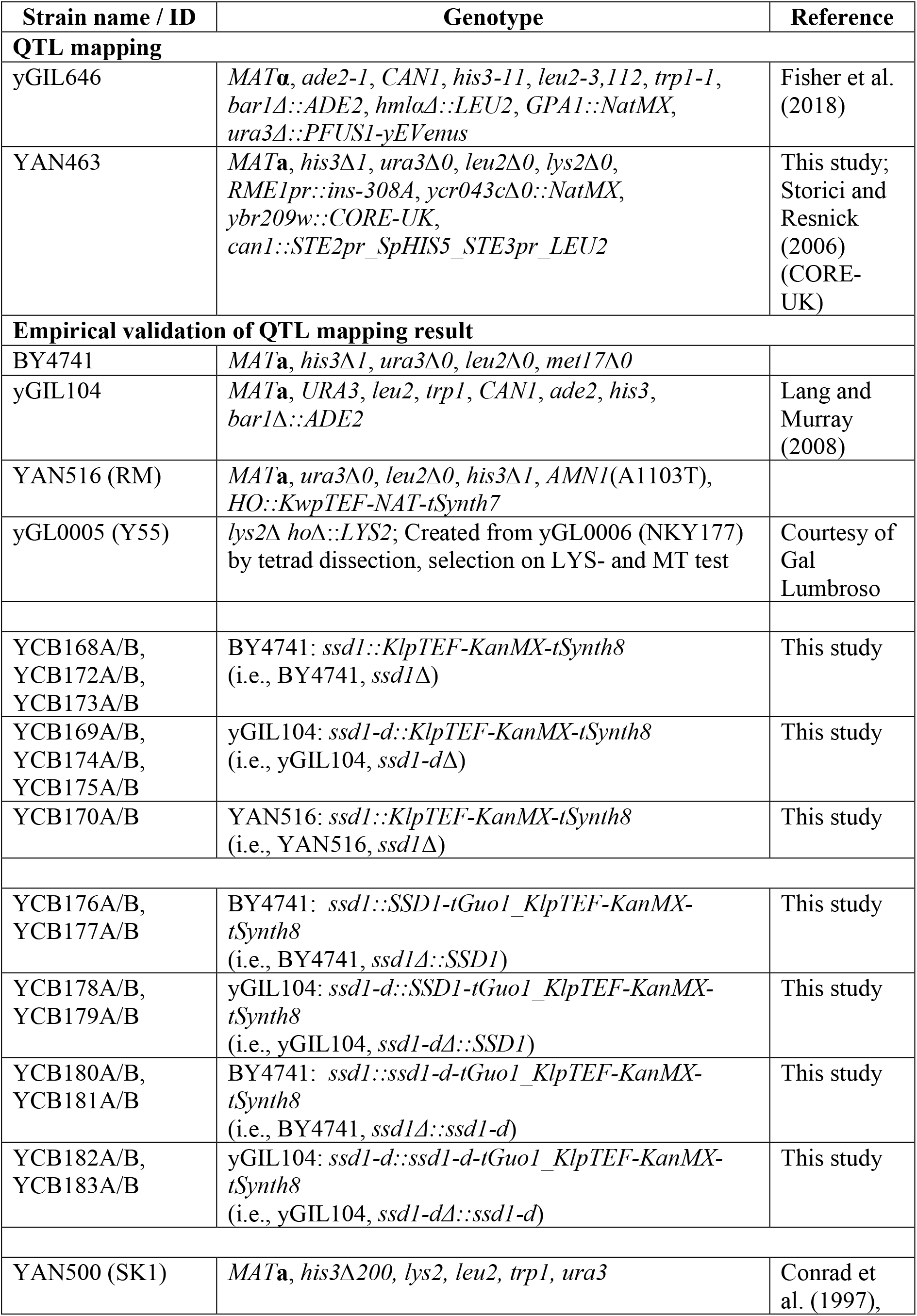
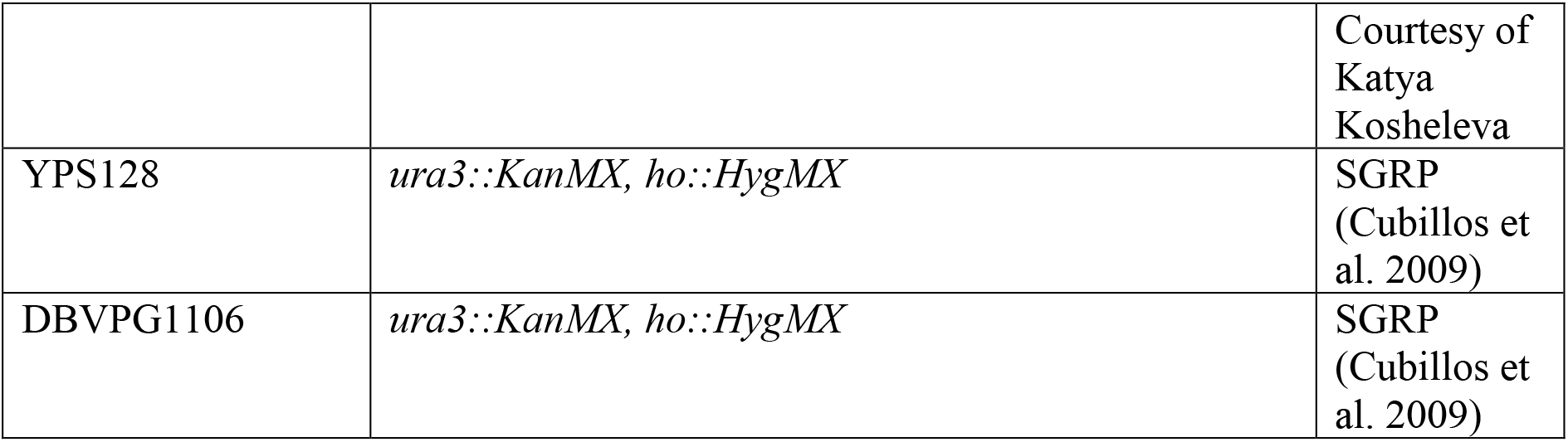
List of experimental strains used and their genotypes

Further, starting with the BY and W303 strains in which HygMX replaced the *SSD1* ORF, we (re-)integrated the BY and W303 *SSD1* alleles alongside KanMX, as described above. The *SSD1* alleles were placed under the strains’ native *SSD1* promoters and terminated by tGuo1, just upstream of KanMX (Curran et al. 2015). This produced versions of BY4741 and W303 in which either the BY or W303 *SSD1* allele was present at the *SSD1* locus (i.e., four strains total). As a control, in the BY and W303 strains in which HygMX was used to knock out *SSD1*, we replaced HygMX with KanMX, producing a set of KanMX-based *SSD1* knockouts ostensibly identical to those described above.

Yeast transformations for strain construction were conducted as described by Gietz (2015), introducing new genetic material as PCR amplicons for incorporation by homologous recombination. A list of the primers used is provided in Table 2. Colony PCR and Sanger sequencing was used to confirm proper integration of amplicons. During strain construction, independent transformant colonies were picked at each step to produce biological replicates and mitigate the phenotypic effects of any unintended off-target mutations. Sytox staining and flow cytometry were used to verify that all ancestral strains were indeed haploid.

**Table 2:**
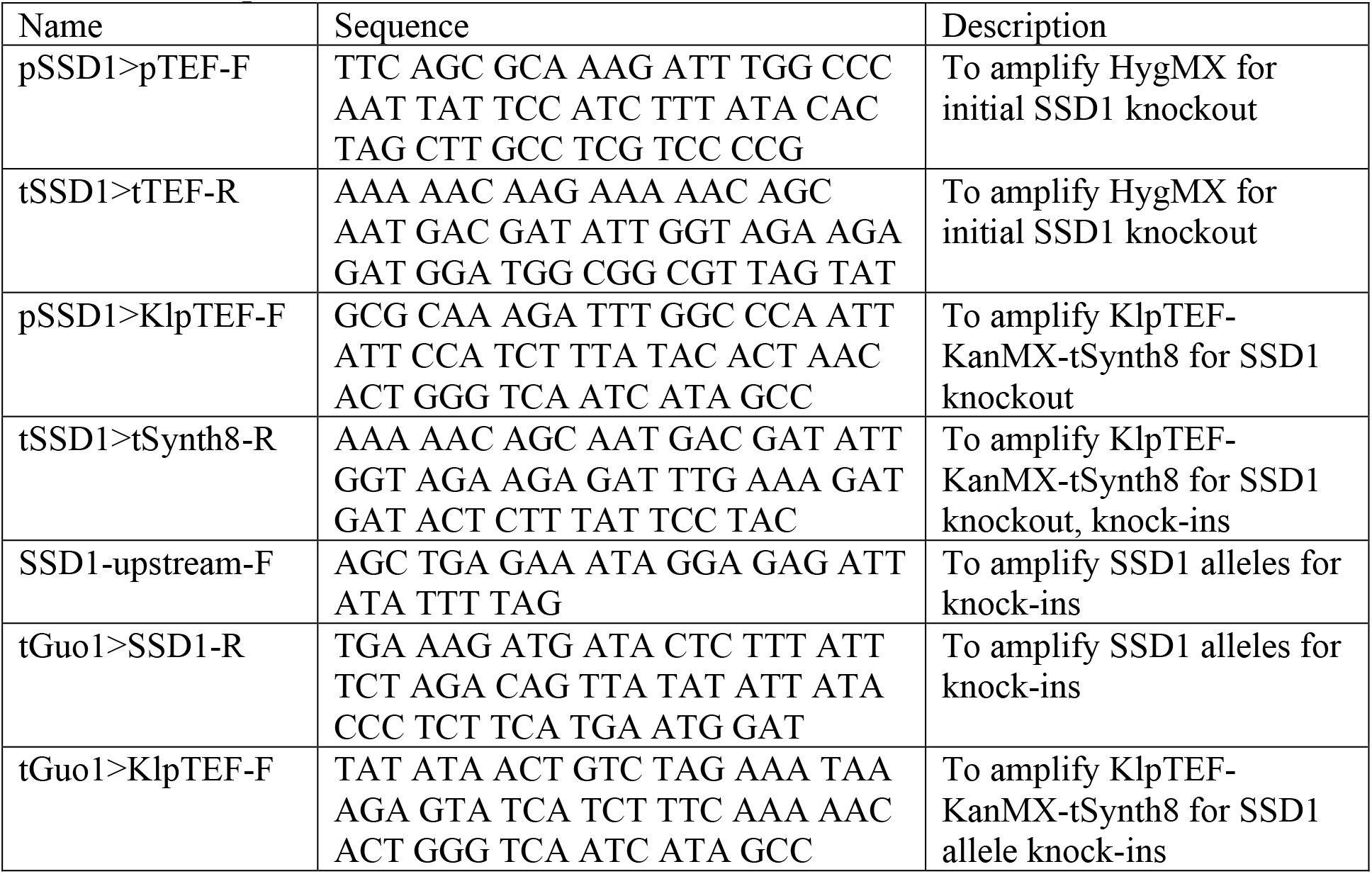
List of primers used in strain construction

We compared the tendency for populations founded with these strains to autodiploidize with each other and with corresponding parental controls by clonally propagating them for 500 generations alongside parental controls and examining their ploidy status after evolution by Sytox staining and flow cytometry. There were 22 technical replicates for each strain construct except for BY4741, *ssd1*Δ and yGIL104, *ssd1-d*Δ, which had 44 technical replicates each. One well for yGIL104, *ssd1-d*Δ::*SSD1* was contaminated by bacteria and thereafter removed. Technical replicates of each genotype were split among at least two biological replicates of that genotype. Populations were frozen initially and at 50-generation intervals in 8% glycerol.

Additionally, we investigated autodiploidization propensity of two domesticated (SK1, Y55) and two wild *S. cerevisiae* strains (YPS128, DBVPG1106) following 500 generations of evolution, using a similar approach to the above with at least twelve technical replicates each. All these strains harbor a functional *SSD1* gene. A consolidated list of all the strains and their genotypes used in this study is provided in Table 1.

### Data availability

Data used for all the figures are available in the SOM file 2. All the strains used here are available from the corresponding author upon request. Raw DNA sequencing reads have been deposited in the NCBI BioProject database with accession number PRJNA713332.

## RESULTS

### Autodiploidization propensity differs across two closely related laboratory strains of *Saccharomyces cerevisiae*

To investigate the intrinsic difference in autodiploidization between BY and W303 populations, we founded seven populations from single clones of each the BY-derived YAN463 and W303-derived yGIL646, respectively, and evolved these for 500 generations. After evolution, we found that all seven replicate YAN463 populations and none of the yGIL646 populations fixed autodiploids (Figure 1A and Figure S2). We also found that seven replicate populations founded by the specific ancestral isolate of yGIL646 used in Fisher et al. (2018) also failed to fix autodiploids during 500 generations of evolution.

**Figure 1.**
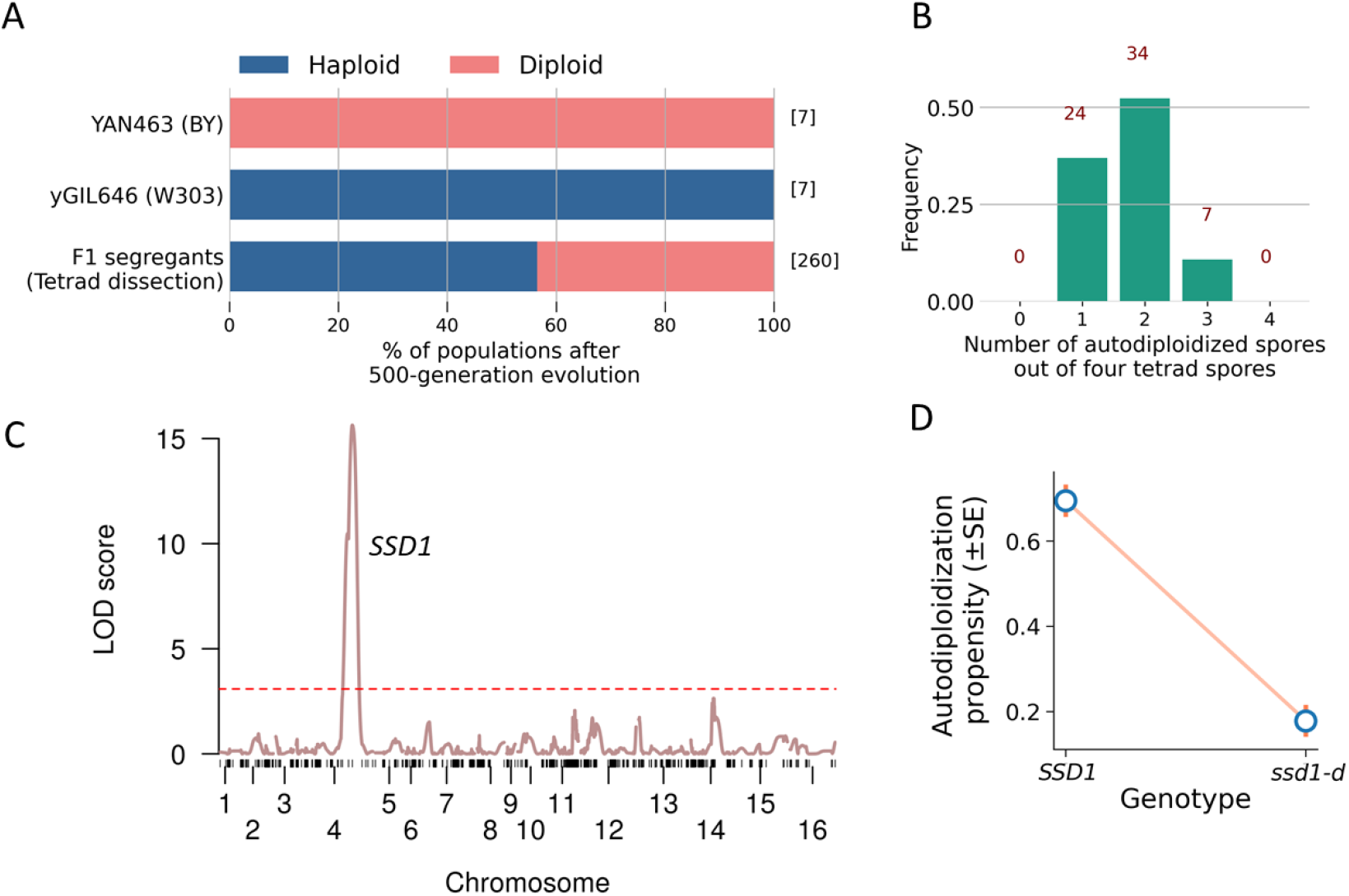
QTL mapping identified a single locus driving variation in autodiploidization propensity. **(A)** Percentage of populations autodiploidized among the clonal replicates of the two parental strains (YAN463 and yGIL646) and their F1 segregants (tetrad spores) after evolving for 500 generations. The numbers inside square brackets denote the number of populations in each category. **(B)** Histogram of the number of autodiploidized spores out of four spores in a tetrad. The numbers in red denote the number of tetrads in each category. **(C)** LOD score for variation in autodiploidization is plotted against the genetic map. The red dashed line indicates a 5% LOD significance threshold computed from 10,000 permutations. The one statistically significant QTL contains a single SNP in the *SSD1* gene. **(D)** Autodiploidization propensity conditional on BY (*SSD1*) and W303 (*ssd1-d*) alleles respectively across all tetrad spores.

In parallel, we conducted a QTL evolution experiment (Figure S1). We first crossed and sporulated yGIL646 and YAN463, dissecting 65 tetrads to obtain 260 F1 segregants. We then founded one population from each of these segregants, and evolved in rich media at 30°C in 96-well plates for 500 generations. Close to half of these populations autodiploidized within 500 generations (44%, 113 out of 260 spores; Figure 1A). In 52% of the tetrads (34/65), two out of four spore-derived populations diploidized while the other two remained haploid. In 37% and 11% of the tetrads, one and three spore-derived populations diploidized, respectively. In none of these tetrads did all four spore-derived populations autodiploidize or remain haploid (Figure 1B).

While the tetrad spores were well-suited to allow mapping of strong QTLs, we predicted that QTL inference might be hindered if the various combinations of auxotrophic markers, drug markers, and mating types in these spores affected autodiploidization. To hedge against this possibility, we also evolved 367 clonal *MAT***a** populations founded by unique “selected spores” from the same cross, bearing one of two sets of common auxotrophies (see Methods). Among these, 184 populations autodiploidized and 179 remained haploid, with 4 ambiguous (Figure 2A).

**Figure 2.**
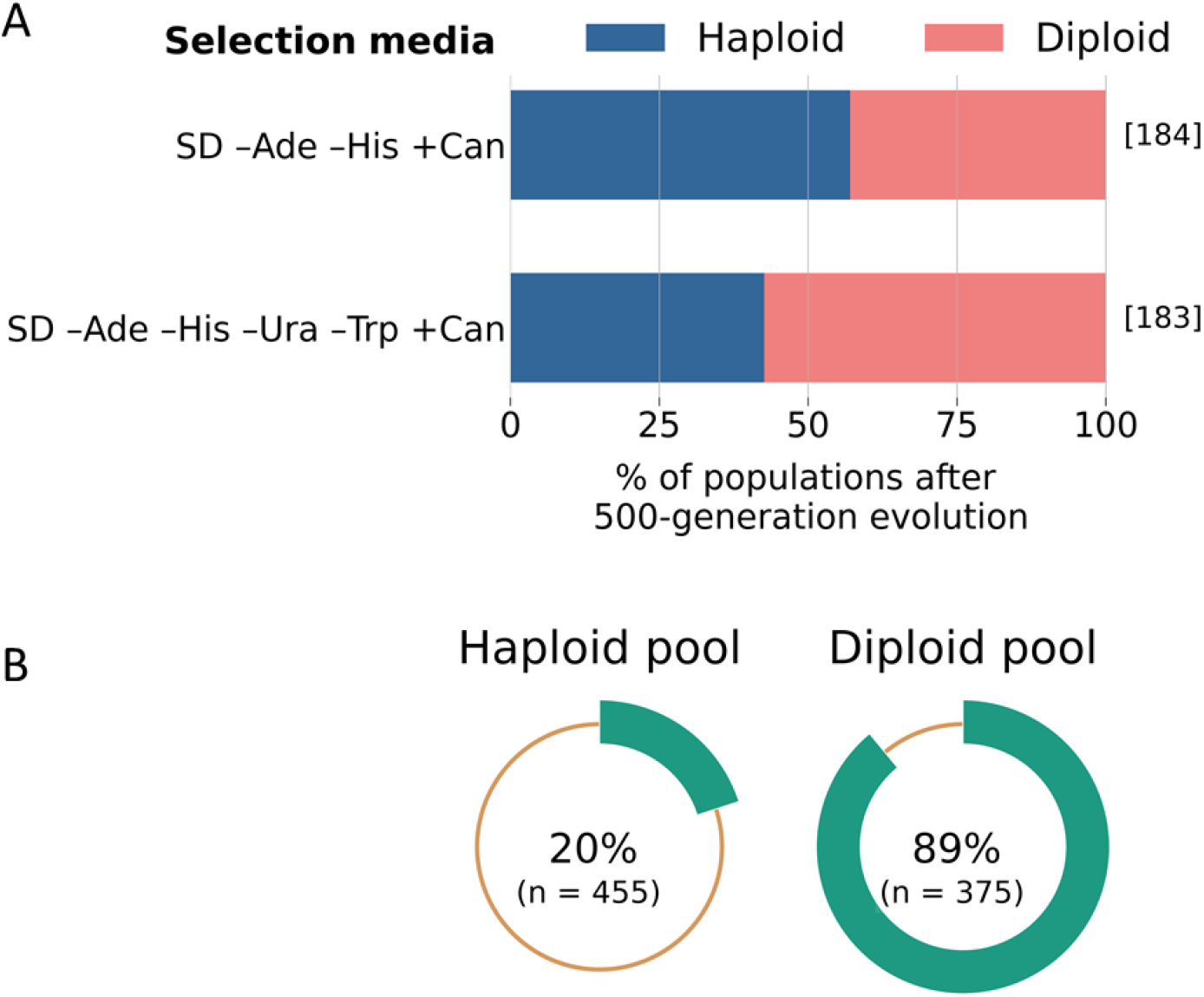
Ploidy status of the ‘selected spores’ after evolution, and enrichment of the BY allele of *SSD1* in diploids. **(A)** Percentage of populations autodiploidized among the spores selected in SD –Ade –His +Can and SD –Ade –His –Ura –Trp +Can media after evolving for 500 generations. The numbers inside square brackets denote the number of populations in each category. Populations with ambiguous ploidy status are shown as haploids. **(B)** Percentage of sequencing reads at *SSD1* locus matching BY allele in haploid and diploid pools of the ‘selected spores.’ Here n denotes the total number of reads at *SSD1* locus for each pool.

### *SSD1* drives differential autodiploidization propensity

To investigate the genetic basis of the difference in autodiploidization propensity between YAN463 and yGIL646, we sequenced each of the 260 F1 segregants in the tetrad set. We then conducted a standard QTL mapping analysis to identify associations between each SNP in the cross and the phenotype described above (specifically, whether the population founded by that segregant autodiploidized after 500 generations of laboratory evolution). We found a single strong QTL on chromosome IV (Figure 1C, LOD = 15.64, *p* < 0.004). The second highest LOD score belonged to a locus on chromosome XIV, but this was not statistically significant (LOD = 2.64, *p* = 0.13). These results remained unaltered when the above analysis was instead performed using the Haley–Knott regression method (Figure S4).

To further evaluate whether any other QTLs played a significant role in determining this phenotype, we performed a test for multiple QTLs that allowed for interactions between loci. Using ~58% of our populations as a test set, we employed a forward/backward stepwise search algorithm to develop a model that allowed for up to 4 interacting QTLs (see Methods for details). However, this search process ultimately found that a single-QTL model implicating the same chromosome IV locus performed best. This model also fit the held-out data (χ^2^ test, *p* < 10^-9^, F test, *p* < 10^-9^), yielding an overall LOD score of 8.63 and explaining 31.5% of the variance in the data.

To confirm this result, we performed a separate single-QTL analysis on the original dataset in which we regressed out the chromosome IV QTL. This analysis yielded no additional statistically significant QTLs (Figure S5).

We found that the BY-allele of the chromosome IV QTL conferred a higher autodiploidization propensity (mean effect ± SE= 0.69 ± 0.04), while the W303-allele diminished autodiploidization in evolving populations (mean effect ± SE = 0.18 ± 0.04; Figure 1D).

To identify the specific gene underlying the significant chromosome IV QTL, we performed nucleotide BLAST (Madden 2013). This algorithm uniquely mapped the QTL to a single SNP in the *SSD1* gene. In BY, *SSD1* codes for a 1250aa-long mRNA-binding translational repressor. By contrast, the W303 *SSD1* allele (henceforth *ssd1-d*) harbors a G →C substitution resulting in a premature stop codon at the ORF’s 698^th^ codon (Y698*). This nonsense mutation effectively truncates the ORF by ~44%.

To verify the findings of the tetrad experiment, we grouped the ‘selected spores’ based on their ploidy and auxotrophy status (see Methods for details) and obtained metagenomic sequences of those pooled samples. Analyzing this data, we found that the proportion of reads matching the BY allele (*SSD1*) was substantially lower in haploid pools than in diploid pools (Figure 2B), irrespective of their auxotrophic status (Figure S6). These results provide independent evidence that *SSD1* is the primary determinant of divergent autodiploidization propensity in clonal BY and W303 populations.

### Populations with a functional copy of *SSD1* autodiploidize more frequently

To test the findings of the QTL mapping analysis described above, we used variants of HygMX and KanMX cassettes (see *Methods*) to construct BY4741 (BY) and yGIL104 (W303) strains in which their *SSD1* alleles had been either swapped or knocked out entirely, with appropriate controls. In total, we produced 3 strains on the BY background (BY4741, *ssd1*Δ; BY4741, *ssd1*Δ::*SSD1*; and BY4741, *ssd1*Δ:: *ssd1-d*) and 3 on the W303 background (yGIL104, *ssd1*Δ; yGIL104, *ssd1*Δ::*SSD1*; and yGIL104, *ssd1*Δ::*ssd1-d*). Biological replicates of each strain were produced during the cloning process. Allele swaps were generated by knocking out *SSD1* with HygMX and re-introducing the appropriate allele with KanMX. Knockout strains were constructed by directly transforming KanMX into the *SSD1* locus or, as a control, by using KanMX to replace HygMX in the penultimate strains in the allele swap constructions.

We founded at least 22 haploid populations from each of these genotypes, divided among the available biological replicates. As in the previously described evolution experiment, we propagated these populations in rich media supplemented with ampicillin on 24-hour cycles, diluting 1024-fold each day and freezing portions of each population every 5 days.

As before, we found that almost all populations of the BY strain bearing its native *SSD1* allele autodiploidized during evolution in rich media for 500 generations, while most populations founded by the W303 strain remained haploid. However, populations founded by either BY or W303 strains in which SSD1 had been knocked out mostly remain haploid (Figure 3). Similarly, populations founded by BY and W303 strains in which the native *SSD1* allele was replaced by the W303 *ssd1-d* allele also mostly remain haploid. In contrast, populations founded by BY or W303 strains in which the native *SSD1* allele was replaced by the BY version of *SSD1* mostly autodiploidize over the course of 500 generations of evolution (Figure 3).

**Figure 3.**
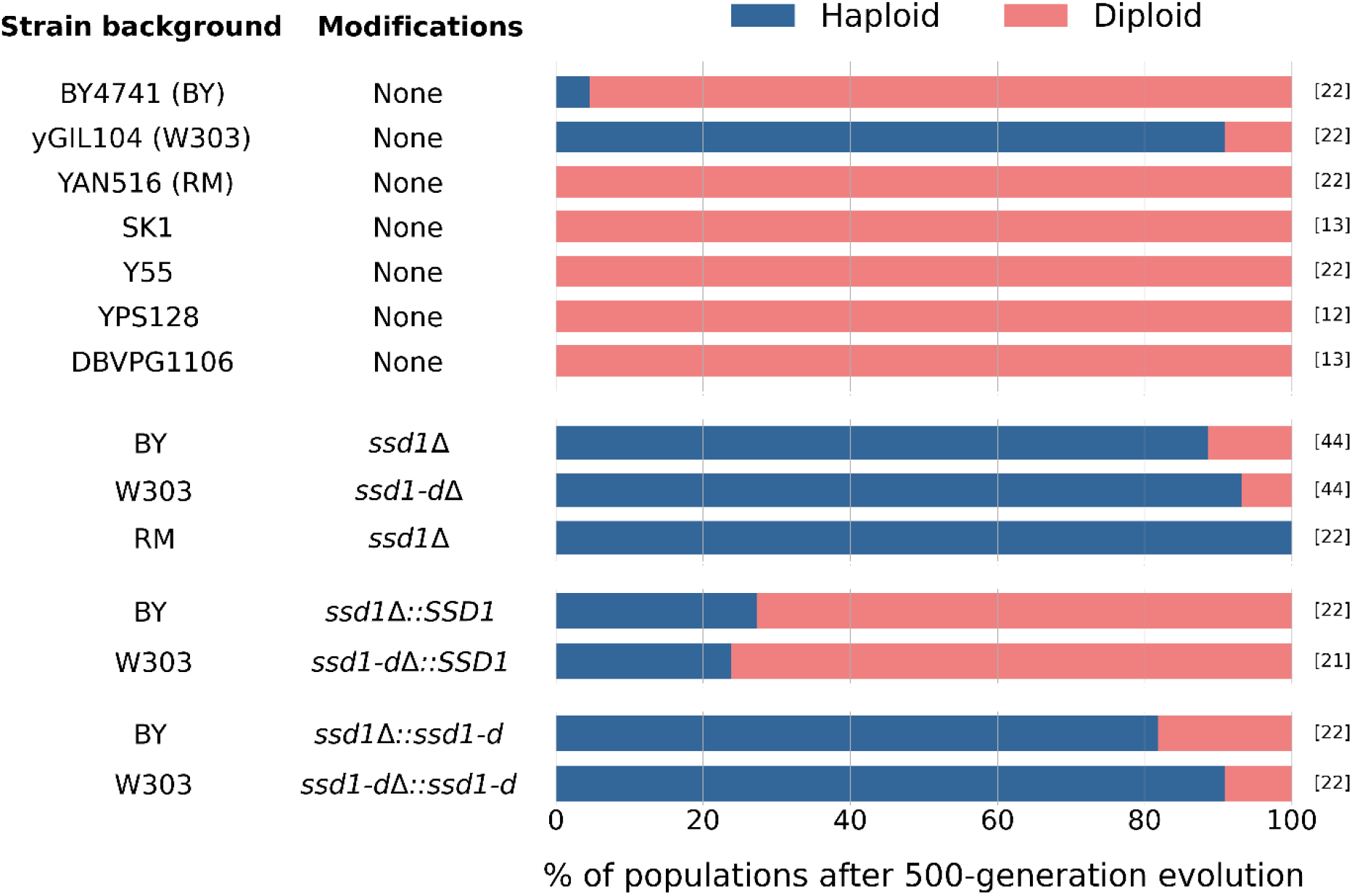
The effect of *SSD1* on autodiploidization. A non-functional *SSD1* gene reduced autodiploidization in W303 populations, while BY, RM, and other domesticated and wild strains expressing full length Ssd1 protein autodiploidized with high frequency. Knocking out *SSD1* reduced autodiploidization in BY and RM, making their frequency similar to that of W303. Allele swap experiments showed that irrespective of the genetic background, presence of the allele expressing the full length Ssd1 protein led to increased autodiploidization, whereas the allele expressing truncated Ssd1 protein reduced it. The numbers in square brackets denote the total number of clonal replicates for each strain. The full genotype of each strain can be found in Table 1.

To evaluate whether these results generalized to more distantly related *S. cerevisiae* strains, we also evolved 12 to 22 replicate populations founded by five other yeast strains (RM11-1a (RM), SK1, Y55, YPS128, and DBVPG1106) for 500 generations. These strains all contain functional copies of *SSD1*, similar to BY. We find that all evolved populations diploidized over the course of evolution (Figure 3). To understand whether *SSD1* played an important role in this phenotype for other strains, we constructed versions of RM in which the native *SSD1* was knocked out with KanMX, just as it was in BY and W303. We evolved 22 replicate populations founded with this knockout genotype (spread across two biological replicates) for 500 generations. We found that knocking out *SSD1* prevented autodiploidization in all replicates (Figure 3).

## DISCUSSION

Ploidy changes mark a major shift in the biology of an organism, with potential consequences for the evolutionary dynamics of populations in which they occur. Although such ploidy changes have been seen frequently in natural, laboratory, and clinical settings, the genetic and environmental factors that influence these changes remain largely unknown. In this study, through experimental evolution and QTL mapping analysis, we find that the gene *SSD1* plays a central role in the emergence and fixation of diploids through spontaneous whole-genome duplication in evolving haploid yeast populations. Our results show that a fully functional *SSD1* gene promotes population autodiploidization, whereas a complete knockout or hypomorphic variant of this gene (as observed in 7 of ~1,000 sequenced isolates (Peter et al. 2018, Scopel et al. 2020)) impedes it substantially.

Further work is needed to understand exactly how *SSD1* affects autodiploidization during experimental evolution. The Ssd1 protein is known to affect many important traits, such as aging, responses to stress, cell wall integrity, and bud formation (Kurischko et al. 2011; Li et al. 2013; Kaeberlein and Guarente 2002; Kaeberlein et al. 2004; Hu et al. 2018; Miles et al. 2019). This pleiotropic footprint makes it hard to speculate about the ultimate mechanisms responsible for *SSD1*’s effect on autodiploidization. For example, one recent study implicated *SSD1* in the maintenance of regular mitochondrial physiology and cytosolic proteostasis crucial for aneuploidy tolerance in wild yeast, showing that that W303 is sensitive to aneuploidy toxicity, which can be rescued with a functional copy of *SSD1* (Hose et al. 2020). Other recent work also provides evidence that yeast lacking *SSD1* are less tolerant of aneuploidies, and it seems this deficiency can be complemented by provision of either of two common functional *SSD1* alleles (Scopel et al. 2020). A similar mechanism may lead to reduced fitness for autodiploidized W303 cells as well, precluding their proliferation in the population. Additionally, previous studies have shown that cell volume roughly doubles with doubling ploidy (Storchova 2014 and references therein). This may make proper *SSD1* function more critical in diploids than haploids, as it is a key regulator of cell wall growth and remodeling. Moreover, another recent study of budding yeast showed that *SSD1* facilitates entry, longevity, and recovery from cellular quiescence (Miles et al. 2019). W303 was shown to have diploid-specific defects in cellular quiescence and stationary phase viability that could be rescued by the introduction of a functional *SSD1*.

Together, these pieces of evidence suggest that a lack of functional Ssd1 protein in W303 cells may mediate the observed differences in population autodiploidization propensity by conferring a fitness disadvantage on autodiploids, independent of the frequency with which they occur de novo in the population. Of course, it is possible that *SSD1* also modulates the baseline per-division frequency of autodiploidization, or influences autodiploid fixation by other, more complex mechanisms (Gerstein and Otto 2011). Delineating these mechanisms is beyond the scope of the current study and a ripe area for future work.

In addition, while populations bearing *SSD1* knockouts or *ssd1-2* typically remained haploid over 500 generations of evolution in these experiments, an appreciable proportion did in fact autodiploidize (Figure 3). This suggests that, in addition to the underlying per-division rate of diploidization and the relative fitness of newly minted diploids, dynamical factors such as clonal interference or the shifting distribution of fitness effects may also substantially influence the likelihood of autodiploid fixation. Further, although our findings point to a likely genetic explanation for differing frequencies of autodiploidization historically observed among yeast evolution experiments, it contrasts with the findings of Fisher et al. (2018), who observed autodiploids take over at high rates in adapting haploid W303 populations. Future work will be necessary to resolve this apparent discrepancy.

In conclusion, we have shown that the frequency at which autodiploids take over adapting populations differs substantially between two closely related laboratory strains of *S. cerevisiae*. We have identified *SSD1* as the key genetic factor underlying the reduced autodiploidization in W303 compared to other strains. Using multiple laboratory and wild strains of *S. cerevisiae*, we showed that, irrespective of genetic background, strains with a functional copy of *SSD1* autodiploidize more frequently, while knocking out or truncating this gene reduces autodiploidization propensity. The results from this study suggest one strategy for modifying the frequency with which diploids take over experimental haploid budding yeast populations. Additionally, we speculate that *SSD1* may be a potential target for modifying the rate of ploidy changes and genome stability in commercial settings, such as the large-scale production of economically important metabolites, and in clinical scenarios, such as the treatment of pathogenic fungal diseases and some cancers.

## Acknowledgments

We thank Katherine Lawrence for helping us with spore genotype inference, Milo Johnson and Shreyas Gopalakrishnan for sharing their high-throughput sequencing library preparation protocol, Gal Lumbroso for providing the strain of Y55 used in this study, Greg Lang for providing the specific ancestral isolate of yGIL646 used in Fisher et al. (2018), and Sean Buskirk for helping in shipping the strain to us. We also thank Andrew W. Murray and Greg Lang for comments on the manuscript. S.T. acknowledges the B4 Science and Technology Fellowship program, funded by DBT, Govt. of India, and OEB department, Harvard University for personal subsistence during this project. C.B. acknowledges the support of the Department of Defense (DoD) through the National Defense Science & Engineering Graduate (NDSEG) Fellowship Program. A.P. acknowledges the support of the Howard Hughes Medical Institute through the Hanna H. Gray Fellows Program. M.M.D. acknowledges support from grant PHY-1914916 from the NSF and grant GM104239 from the NIH. The computations in this paper were run on the FASRC Cannon cluster supported by the FAS Division of Science Research Computing Group at Harvard University.

## Supplementary information

### Text S1. Data clean up prior to QTL analysis

Based on standard recommendations (Broman and Sen 2009), prior to QTL mapping following diagnostic probes were computed to ensure quality and integrity of the dataset.

#### Segregation distortion

Under normal circumstances BY and W303 alleles for each locus should segregate equally. To test this, we inspected genotype frequencies at each marker locus using function `geno.table`. 30 loci failed χ2 test for deviation from Mendelian proportions (i.e. 1:1, here). They were dropped from subsequent analysis.

#### Compare individuals’ genotypes

In order to identify pairs of segregants with unusually similar genotypes across all loci, we compared genotypes for each pair of individuals using the `comparegeno` function. One pair of segregants had >99% similarity in genotype identity, was detected as an outlier (Grubb’s test: Q = 5.81, *p* = 0.0002) and therefore removed from the subsequent analysis.

#### Counting crossovers

The number of crossover events observed for each segregant was computed using the `countXO` function. The number of crossovers was found to be unreasonably high for one segregant (Grubb’s test: Q = 9.68,*p* <10^-16^), and this segregant was removed from further analysis.

**Figure S1.**
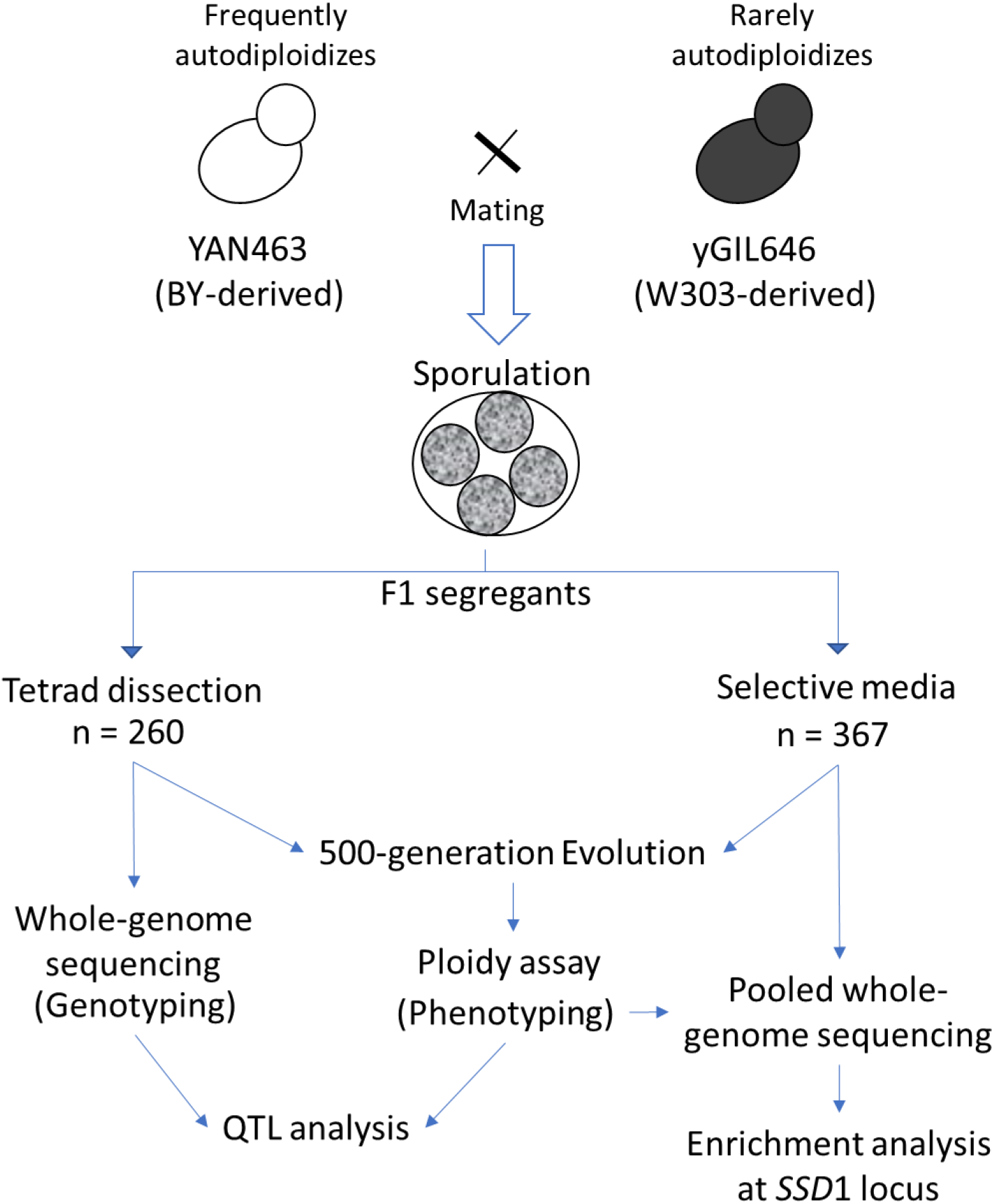
Schematic diagram of the QTL mapping experiment

**Figure S2.**
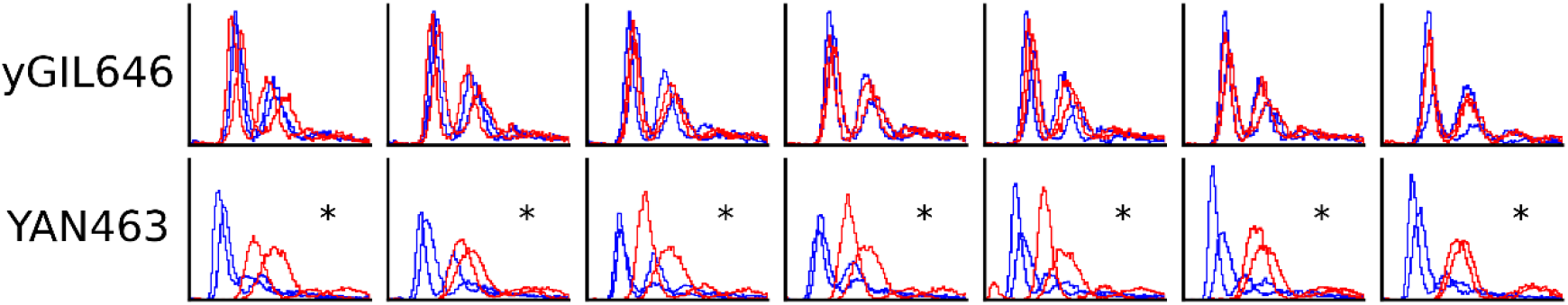
The ploidy state of the 7 replicate populations of the parental strains before and after 500-generation evolution. The plots show FITC histograms of Sytox-stained cells of each population, where the x-axis is in arbitrary fluorescence units (linear), and the y-axis is frequency. Blue and red curves denote the two technical replicate runs for each of the initial and final timepoints of evolution respectively. Populations where autodiploidization has been observed are marked by asterisks.

**Figure S3.**
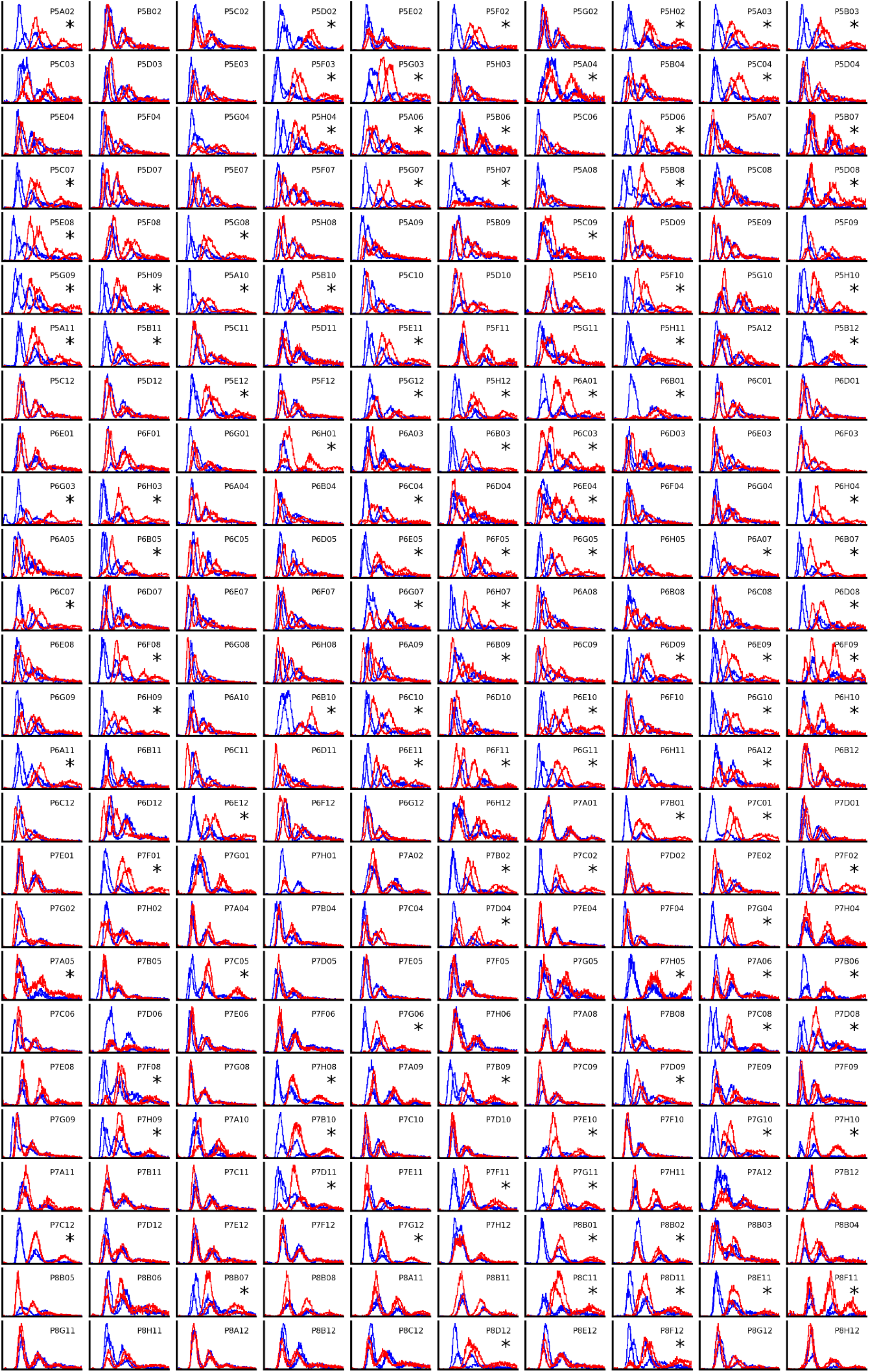
The ploidy state of the 260 tetrad populations before and after 500-generation evolution. The plots show FITC histograms of Sytox-stained cells of each population, where the x-axis is in arbitrary fluorescence units (linear), and the y-axis is frequency. Code starting with ‘P’ on each panel indicates population ID. Blue and red curves denote the two technical replicate runs for each of the initial and final timepoints of evolution respectively. Populations where autodiploidization has been observed are marked by asterisks.

**Figure S4.**
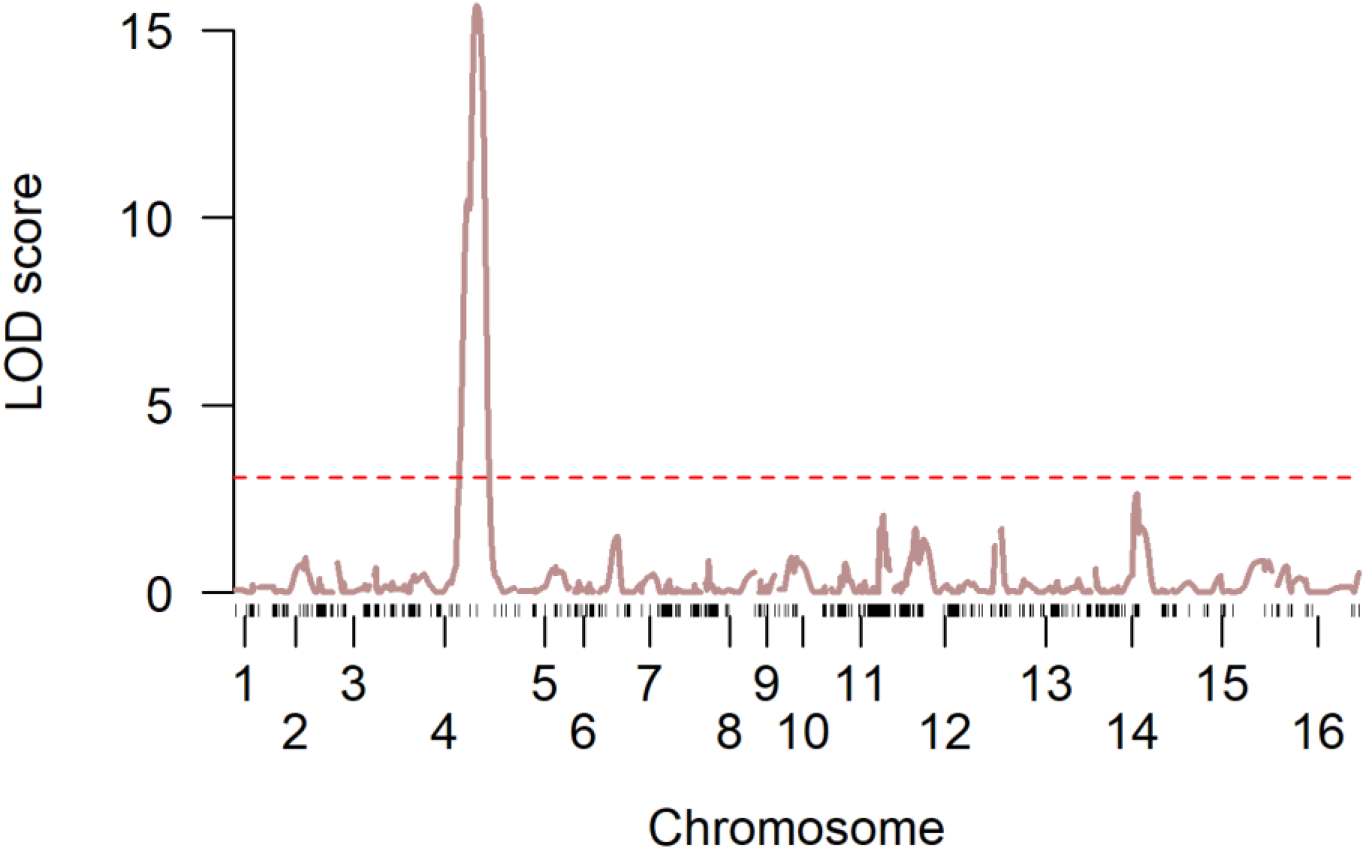
LOD score for variation in autodiploidization obtained using the Haley–Knott regression method is plotted against the genetic map. The red dashed line indicates a 5% LOD significance threshold computed from 10,000 permutations. The single statistically significant QTL is identical to that of Figure 1C and falls within the *SSD1* locus.

**Figure S5.**
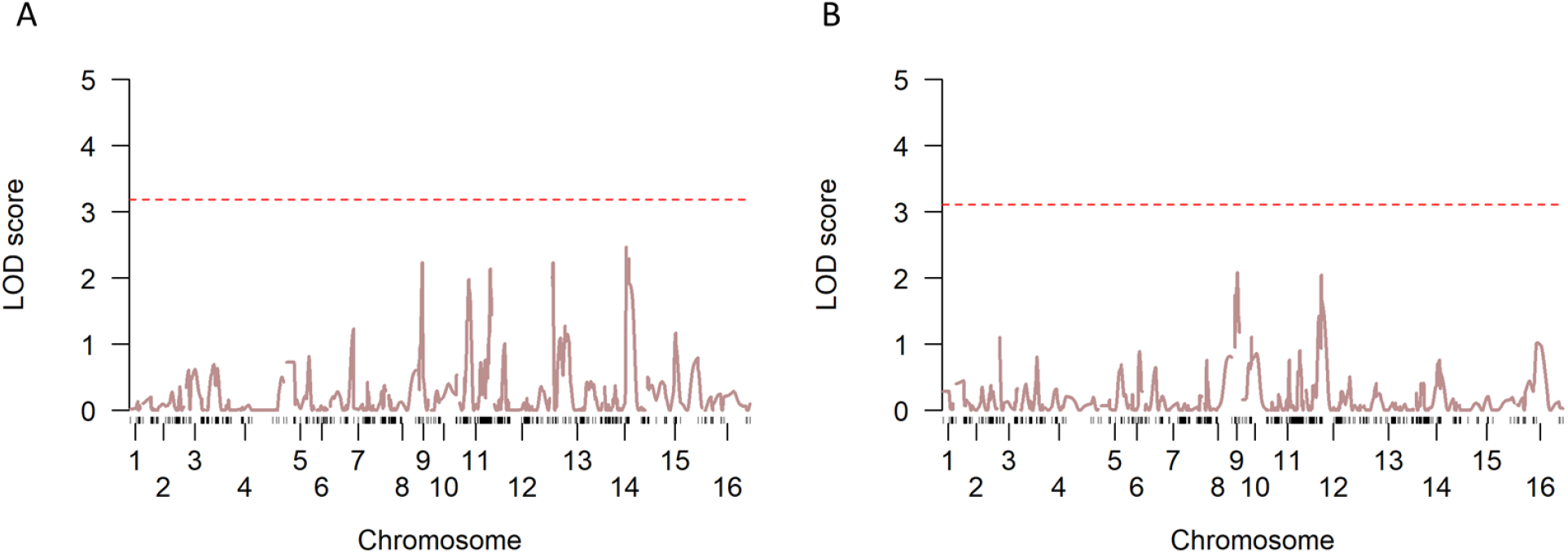
LOD score for variation in autodiploidization obtained using standard interval mapping method after regressing out the statistically significant chromosome IV QTL. The red dashed line indicates a 5% LOD significance threshold computed from 10,000 permutations. No additional statistically significant QTL are present for the segregants with (A) BY and (B) W303 allele of the chromosome IV QTL.

**Figure S6.**
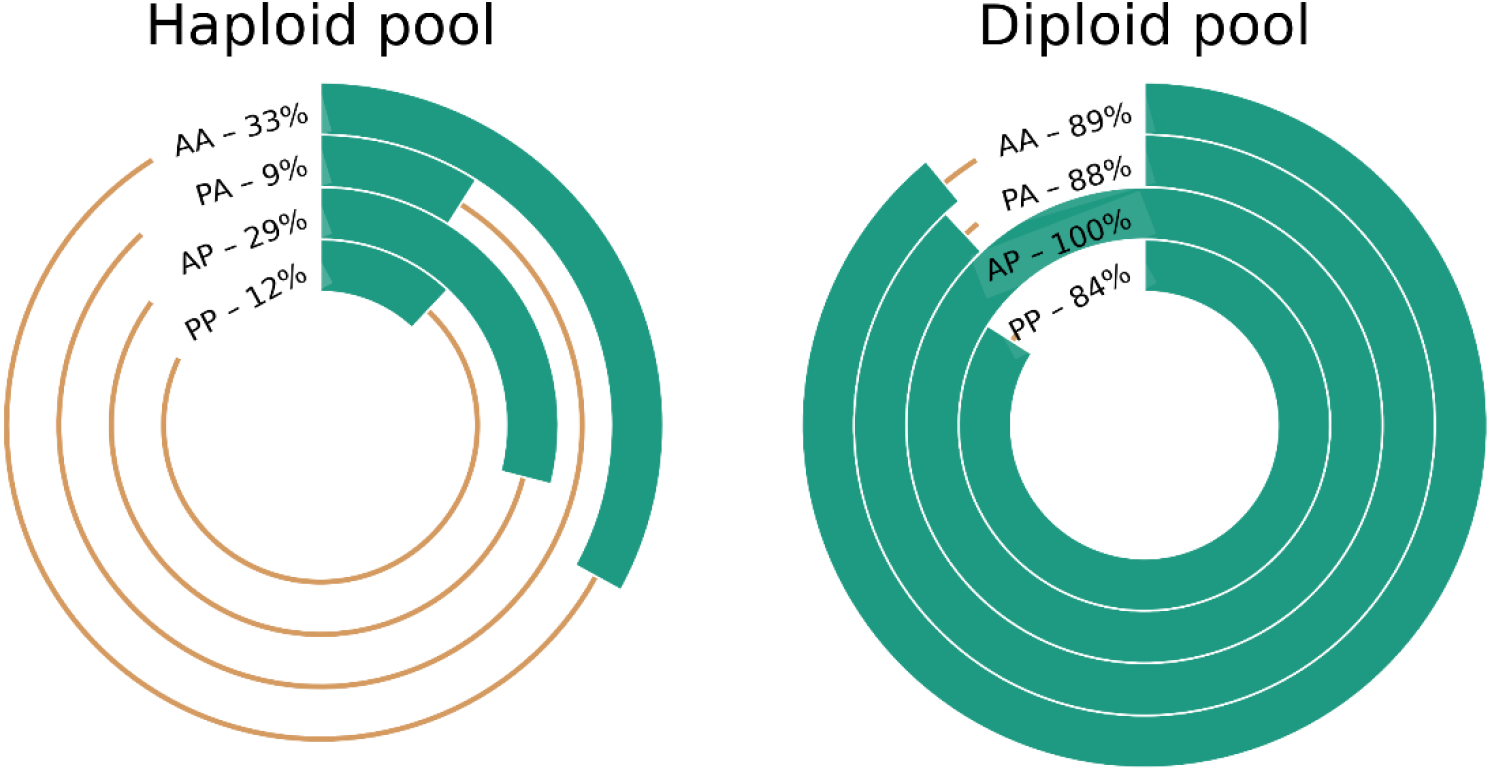
Percentage of sequencing reads at *SSD1* locus matching BY allele in haploid and diploid pools of the ‘selected spores.’ The two-letter code for each plot indicate whether they are auxotrophic (A) or prototrophic (P) for Tryptophan and Lysine, (e.g. ‘PA’ denotes the spores that are prototrophic for Tryptophan but auxotrophic for Lysine). Irrespective of the auxotrophy status, the BY allele is substantially enriched in the diploid pool, whereas it is depleted in the haploid pool.

